# Combined effects of migration distance, foraging method vegetation density, and population density on wing shapes of boreal songbirds

**DOI:** 10.1101/413351

**Authors:** Flavie Noreau, André Desrochers

**Affiliations:** Centre d’étude de la forêt, Université Laval, Pavillon Abitibi-Price, 2405 rue de la Terrasse, Québec, Qc, G1V 0A6 CANADA

**Keywords:** ORNITHOLOGY, COMPARATIVE, ECOLOGY, FLIGHT, LOCOMOTION, MIGRATION, MORPHOLOGY

## Abstract

In birds, migration distance is known to influence morphological attributes that influence flight performance, especially wing shape. However, wing shape is under the likely influence of less documented factors such as foraging method, vegetation density and isolation of individuals and populations. To better understand factors leading to interspecific differences in wing shape, we measured the pointedness of wings (Kipp’s distance) of 1017 live birds of 22 species in an eastern Canadian boreal forest. We modeled wing pointedness as a function of migration distances from eBird records, foraging, habitat, and population density data from *Birds of North America* monographs. Long-distance migrants and species living in low-density vegetation had more pointed wings than shorter-distance migrants and dense-vegetation dwellers, in accordance to our predictions. After accounting for vegetation density and migration distance, we found no link between the extent of aerial foraging or mean breeding population density, an indicator of isolation, and wing pointedness. Those results are consistent with a tradeoff between sustained flight efficiency and maneuverability, but suggest that interspecific variation in wing shape due specifically to foraging method or habitat isolation is nonexistent or obscured by other factors.

## Introduction

Variation in wing shape is constrained by aerodynamic principles, but it remains highly variable from one species to another. In the case of birds, elongated wings with short secondary remiges and long primary remiges facilitate sustained flight (Bowlin and Wikelski 2008) while rounder wings are thought to facilitate maneuverability (Savile 1957). Wing shape is associated to migratory performance in mammals and birds (Palmer 1900; Norberg and Rayner 1987). This association has been shown with Palearctic warblers (Marchetti et al. 1995; Nowakowski et al. 2014), Nearctic thrushes (Dilger 1956) and swallows (Huber et al. 2016), seven Palearctic and Nearctic passerine genera (Mönkkönen 1995), as well as shorebirds (Burns 2003; Minias et al. 2015). North American passerines may provide an exception to this rule (Keast 1980; Niemi 1985), but no quantitative evidence exists to assess this claim.

Once migration is completed, birds are subject to further selective pressure on functional traits related to flight, and at very different spatial scales (Huber et al. 2016). For species gleaning food on the ground, perched or in dense vegetation, the advantages of manoeuvrability may overwhelm the need for efficient sustained flight (Dilger 1956; Niemi 1985). Isolation from conspecifics and concomitant movement may exert evolutionary pressure independent of migration distance, in favor of elongated wings (Desrochers 2010). While their roles are often speculated upon, foraging method, vegetation density and population isolation remain poorly documented correlates of avian wing shape.

In this study we link bird wing pointedness, more specifically the projection of primary flight feathers also known as Kipp’s distance (Lockwood et al. 1998), and four important aspects of their ecology: migration distance, foraging method, breeding habitat density and isolation. With boreal forest songbirds, we test the following hypotheses: wings are more pointed in species 1) with longer migration distances; 2) more frequently using aerial foraging tactics such as hovering and sallying; 3) foraging in sparse vegetation; and 4) found in low-density breeding populations.

## Materials and methods

### Study Area

We conducted field work at Forêt Montmorency, Quebec, Canada (47.4 N, 71.1W) during the summers 2013 and 2014. This forest has been managed for timber since the early 1930’s, with ecosystem management (Bélanger 2001; Gauthier et al. 2008) dominating the southern half of the territory, and large (100-150 ha) clearcuts dominating the northern half. Young stands (<20 y) are generally a dense mixture of deciduous and coniferous species (Mallik et al. 2014), but older stands are largely dominated by balsam fir (*Abies balsamea* (L.) Mill.), with occasional groves of spruce (*Picea glauca* (Moench) Voss, *Picea mariana* (Mill.) Britton), birch (*Betula papyrifera* Marsh.) and poplar (*Populus tremuloïdes* Michx., *Populus balsamifera* L.).

### Wing pointedness

We captured birds in mist nets at 30 sites throughout the 412 km^2^ study area. We deployed nets between 5 am and 11 am over 2 or 3 consecutive days at each site. We used recordings of mobbing calls (Gunn et al. 2000) and species songs to increase capture rates. We banded, identified and measured birds with age and sex determined by plumage (Pyle 1997). To facilitate the measurements of wing pointedness, we took one to three photos of the right wing flattened on a wing ruler for each bird, with a Nikon D80 digital SLR with a resolution of 10 Megabytes in 2013 and Nikon D7100 DSLR 24 megabytes in 2014. We took measurements in pixels from each photo with an image processing software, ImageJ (http://imagej.nih.gov/ij/). Two lengths were obtained: a) the total length of the folded, flattened, wing, and b) the length between the posterior end of the most distal secondary feather and the posterior end of the wing (Fig. 1). Wing pointedness was quantified as the ratio b/(a+b) (Lockwood et al. 1998).

**Fig. 1.**
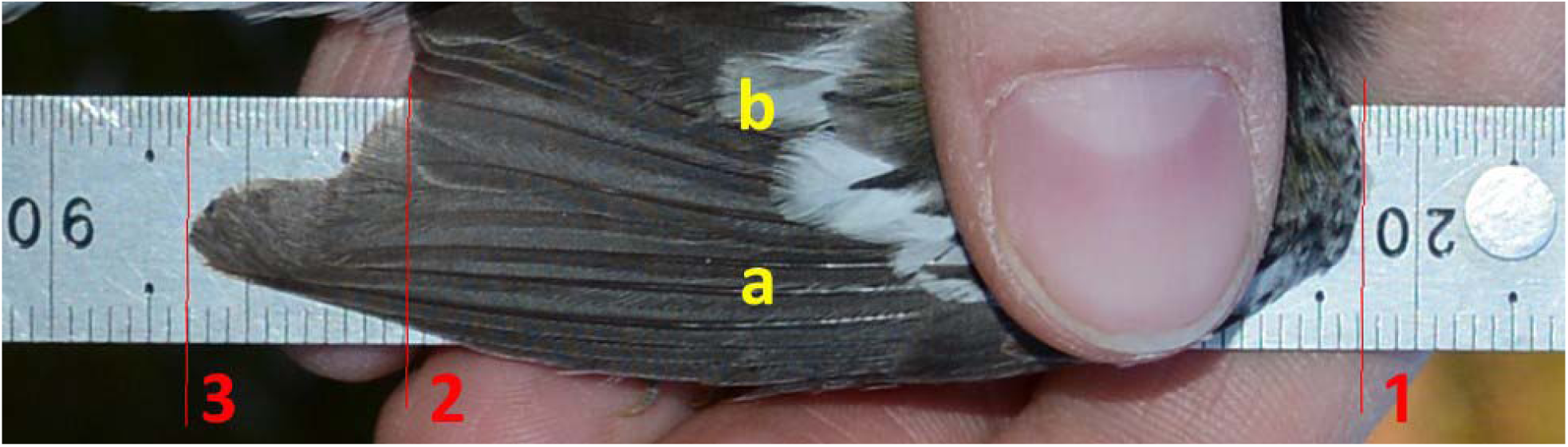
Measurements used to calculate wing pointedness. Red markers (1,2 and 3) were tattooed on photos to facilitate measurements.

Repeated (blind) estimates of the wing pointedness of the same bird by the same author had a mean standard deviation of 0.27 mm (*n* = 666 birds), and a coefficient of variation of 1.6%. The mean standard deviation of wing pointedness estimates of the same bird measured separately by the two authors was 0.3 mm (*n* = 20 birds). We assume that precision and accuracy of wing pointedness estimates were sufficiently high to yield useful interspecific comparisons.

### Aerial Foraging, Vegetation Density, and Population Density

We characterized foraging method, vegetation density, and breeding population density with monographs from *The Birds of North America Online*, BNA (Rodewald 2015). For foraging method, we estimated for each species the proportion of time spent gleaning while perched or on the ground, hovering (picking prey in flight, from substrate), and sallying (flying from perch to catch flying prey, (Eckhardt 1979)). When available, quantitative estimates we used, otherwise we attributed 10% of the time for foraging methods depicted as “occasional” (including “common”, “frequent”, “sometimes”) and 5% for method rarely used (including “seldom”), leaving the rest to the most frequently used method. We calculated an aerial foraging index as a weighted sum, with weights of 1, 2, and 3 for gleaning, hovering and sallying respectively (possible range 1-3; Table 1).

**Table 1.** Species characteristics. Species ordered according to their estimated phylogenetic tree.

| Scientific name | English name | N | Mean Wing pointedness | Aerial foraging index | Vegetation Density index | Mean Migration Distance (km) | Mean Breeding Density (Pairs/km<sup>2</sup>) |
| --- | --- | --- | --- | --- | --- | --- | --- |
| *Setophaga coronata* L. | Yellow-rumped Warbler | 66 | 0.232 | 1.45 | 0.85 | 3526 (74222) | 44.4 |
| *Setophaga virens* Gmelin | Black-throated Green Warbler | 39 | 0.238 | 1.20 | 0.38 | 3771 (2989) | 122.5 |
| *Setophaga ruticilla* L. | American Redstart | 37 | 0.196 | 2.20 | 3.02 | 3554 (4610) | 163.7 |
| *Setophaga striata* Forster | Blackpoll Warbler | 128 | 0.292 | 1.15 | 1.08 | 4836 (230) | 46.1 |
| *Setophaga castanea* Wilson | Bay-breasted Warbler | 26 | 0.283 | 1.15 | 0.97 | 4503 (427) | 155.9 |
| *Setophaga magnolia* Wilson | Magnolia Warbler | 169 | 0.193 | 1.25 | 1.61 | 3759 (2316) | 90.9 |
| *Cardellina pusilla* Wilson | Wilson's Warbler | 27 | 0.190 | 1.35 | 2.72 | 3932 (5636) | 147.8 |
| *Oreothlypis ruficapilla* Wilson | Nashville Warbler | 45 | 0.214 | 1.15 | 0.55 | 3982 (2103) | 56.3 |
| *Oreothlypis peregrina* Wilson | Tennessee Warbler | 18 | 0.265 | 1.15 | 1.82 | 4153 (3023) | 251.2 |
| *Geothlypis trichas* L. | Common Yellowthroat | 12 | 0.161 | 1.15 | 3.14 | 3498 (21240) | 98.0 |
| *Junco hyemalis* L. | Dark-eyed Junco | 49 | 0.197 | 1.00 | 2.97 | 2617 (95691) | 95.8 |
| *Zonotrichia albicollis* Gmelin | White-throated Sparrow | 96 | 0.163 | 1.00 | 3.45 | 2159 (70635) | 47.2 |
| *Passerella iliaca* Merrem | Fox Sparrow | 11 | 0.218 | 1.10 | 2.21 | 2957 (32735) | 95.0 |
| *Melospiza lincolnii* Audubon | Lincoln's Sparrow | 30 | 0.180 | 1.05 | 2.33 | 3703 (16018) | 324.0 |
| *Regulus satrapa* Lichtenstein | Golden-crowned Kinglet | 22 | 0.213 | 1.60 | 2.41 | 2570 (39908) | 169.6 |
| *Regulus calendula* L. | Ruby-crowned Kinglet | 50 | 0.194 | 1.90 | 1.99 | 3398 (68375) | 185.0 |
| *Catharus ustulatus* Nuttall | Swainson's Thrush | 114 | 0.299 | 1.40 | 1.22 | 4930 (1147) | 88.4 |
| *Catharus bicknelli* Ridgway | Bicknell's Thrush | 7 | 0.274 | 1.30 | 2.03 | 3138 (137) | 129.7 |
| *Poecile hudsonicus* Forster | Boreal Chickadee | 13 | 0.159 | 1.20 | 0.00 | 0 (2154) | 6.7 |
| *Vireo philadelphicus* Cassin | Philadelphia Vireo | 24 | 0.237 | 1.15 | 1.83 | 4047 (839) | 38.6 |
| *Empidonax flaviventris* Baird | Yellow-bellied Flycatcher | 14 | 0.217 | 2.42 | 3.45 | 3927 (1042) | 15.4 |
| *Empidonax alnorum* Brewster | Alder Flycatcher | 20 | 0.226 | 1.74 | 3.63 | 6519 (35) | 54.9 |

For vegetation density, we made a systematic search of words qualifying density using the “Habitat” section of the BNA account of each species. The search was divided between breeding and winter ranges. We sought words related to shrub layers, "shrub", "bush", "scrub", "dense", "thick", and measured the location of first occurrence of one of these words (*n*th word in the Habitat section). The relative position of the word in the text (100% = first, 0%=last) was taken as an indicator of weight. We assume that keywords used earlier in the paragraph provided evidence for a stronger link to dense cover than those listed towards the end of the text. We added ratings computed for the breeding as well as the wintering season to obtain a vegetation density index for each species (Table 1).

We calculated the mean breeding population density of each species from all the breeding density estimates cited in the BNA species accounts (Table 1). In the case of the little-known Bicknell’s Thrush (*Catharus bicknelli*) we used Rimmer et al. (1996).

### Migration distance

To assign average species migration distances, we used the eBird database (Sullivan et al. 2009). This site was developed by the Cornell Lab of Ornithology and the National Audubon Society and contains millions of bird sightings throughout the world. We obtained coordinates, rounded to the nearest degree of latitude and longitude, of each observation in January and February (i.e. wintering) for all bird species considered in this study, as well as the total observation effort in minutes. Multiple records of a species in the same latitude/longitude on the same year were reduced to one to prevent bias due to repeated observations (e.g. stragglers attracting large numbers of birders). For each eBird entry (*n =* 445,512, Table 1), we calculated the great-circle distance from the geographic center of the Forêt Montmorency using the ‘sp’ package of R software (Pebesma and Bivand 2005; Bivand et al. 2013).

### Statistical analysis

Studies comparing mean attributes from multiple species may suffer from a lack of true replication because of phylogenetic proximity among subsets of the species. To account for this, we obtained phylogenetic trees of the species studied, from Jetz et al. (2012). Because of uncertainty in the relationship between DNA data and years since speciation, phylogenetic trees are only approximations based on various assumptions about the rate of phylogenetic divergence. Thus, we generated 10,000 phylogenetic trees (Fig. 2) and established phylogenetic distances for each dyad from the 22 study species (Paradis et al. 2004).

**Fig. 2.**
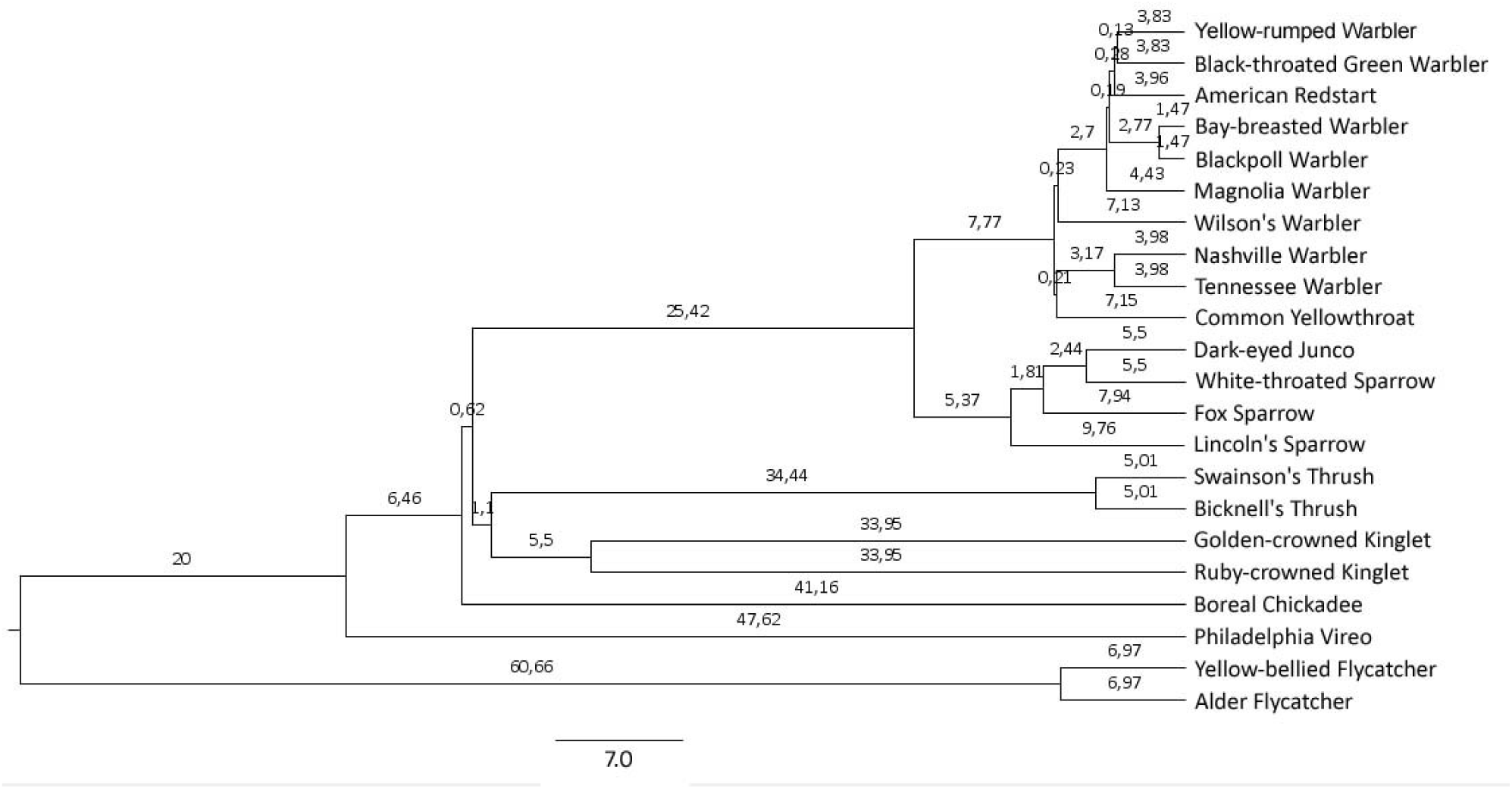
A phylogenetic tree of the species in the study. Generated from the website "birdtree.org" (Jetz et al. 2014).

We modeled the independent influences of migration distance, aerial foraging index, vegetation density index, and population density on wing pointedness with a linear model using phylogenetic Generalized Least Squares (pGLS). The pGLS method accounts for the phylogenetic distance between species by using a distance covariance matrix which gives more weight to differences in wing pointedness between remote species phylogenetically than in wing pointedness of closely related species (Felsenstein 1985; Grafen 1989; Harvey and Pagel 1991). We ran 10,000 pGLS models, corresponding to each of the 10,000 phylogenetic trees of all species. We calculated summary statistics from those 10,000 models for regression coefficients, their standard error, and *p* values. Data analyses were conducted in R, using the ‘ape’ (Paradis et al. 2004), ‘nlme’ (Pinheiro et al. 2016), and ‘phylolm’ (Ho and Ané 2014).

## Results

We considered 22 different bird species, that is, all species with at least five specimens captured (*n* = 1017; Table 1). Intraspecific variance of wing pointedness was significantly lower than interspecific variance (F_21,995_ = 266.2; *p* < 0.001), thus we assume that mean estimates for each species were sufficiently contrasted for comparative analysis. Vegetation density and aerial foraging indices were not significantly, correlated (*r*_*s*_ = 0.1, *p* = 0.6, *n* = 22).

Wing pointedness was positively associated with migration distance (Fig. 3) and negatively associated to vegetation density (Fig. 4) with all phylogenetic trees (Table 2). Results were similar when the ‘outlier’ species with shortest and longest migration distances (Fig. 3) were removed. We found no relationship between wing pointedness, aerial foraging or breeding population density (Table 2) Results obtained with a multiple regression ignoring phylogenetic effects were highly similar for migration distance (estimate = 2.20e-5 ± 5.9e-6, *p* < 0.001), aerial foraging (estimate = 8.87e-4 ± 2.0e-2, *p* = 0.9), vegetation density (estimate = -1.87e-2 ± 7.1e-3, *p* = 0.02), and breeding population density (estimate = -2.38e-4 ± 9.1e-3, *p* > 0.9).

**Fig. 3.**
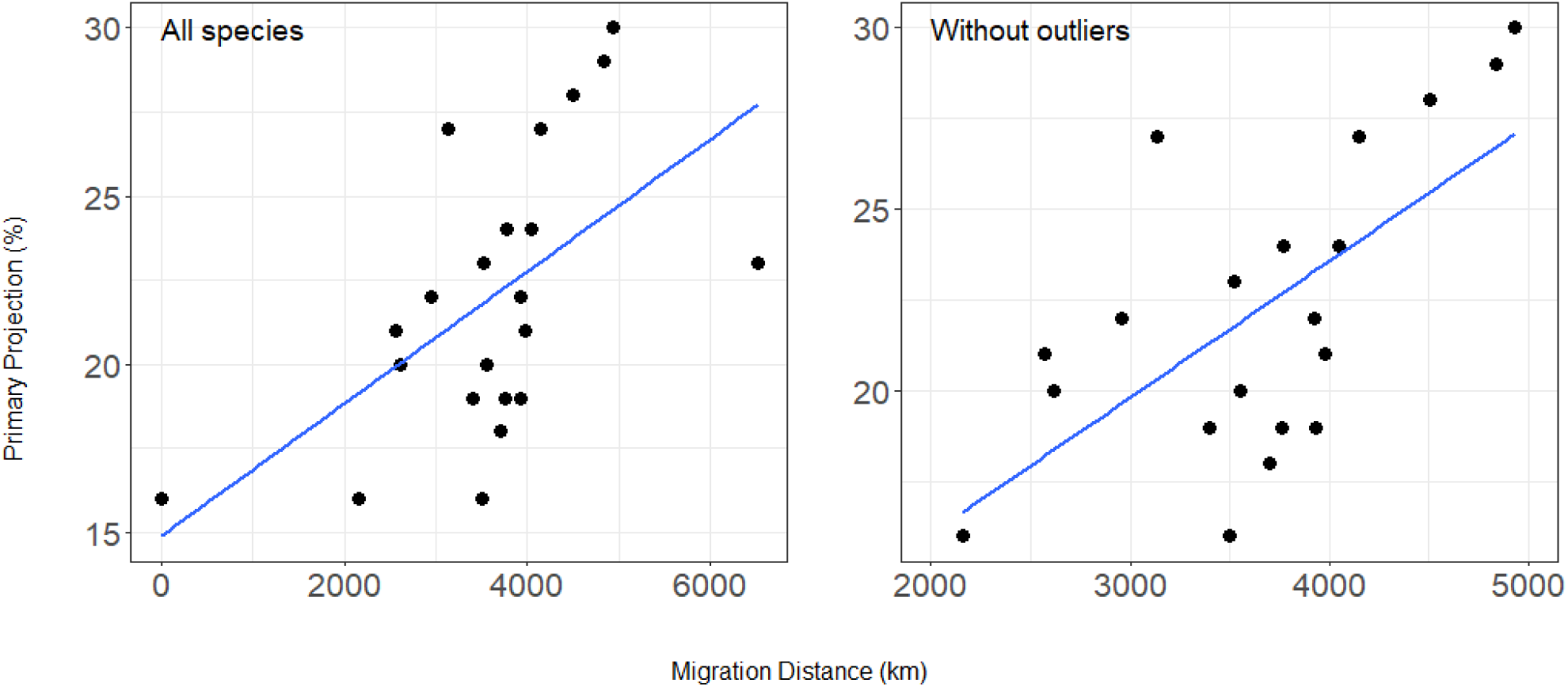
Wing pointedness in relation with migration distance in kilometers. Each point denotes a species.

**Fig. 4.**
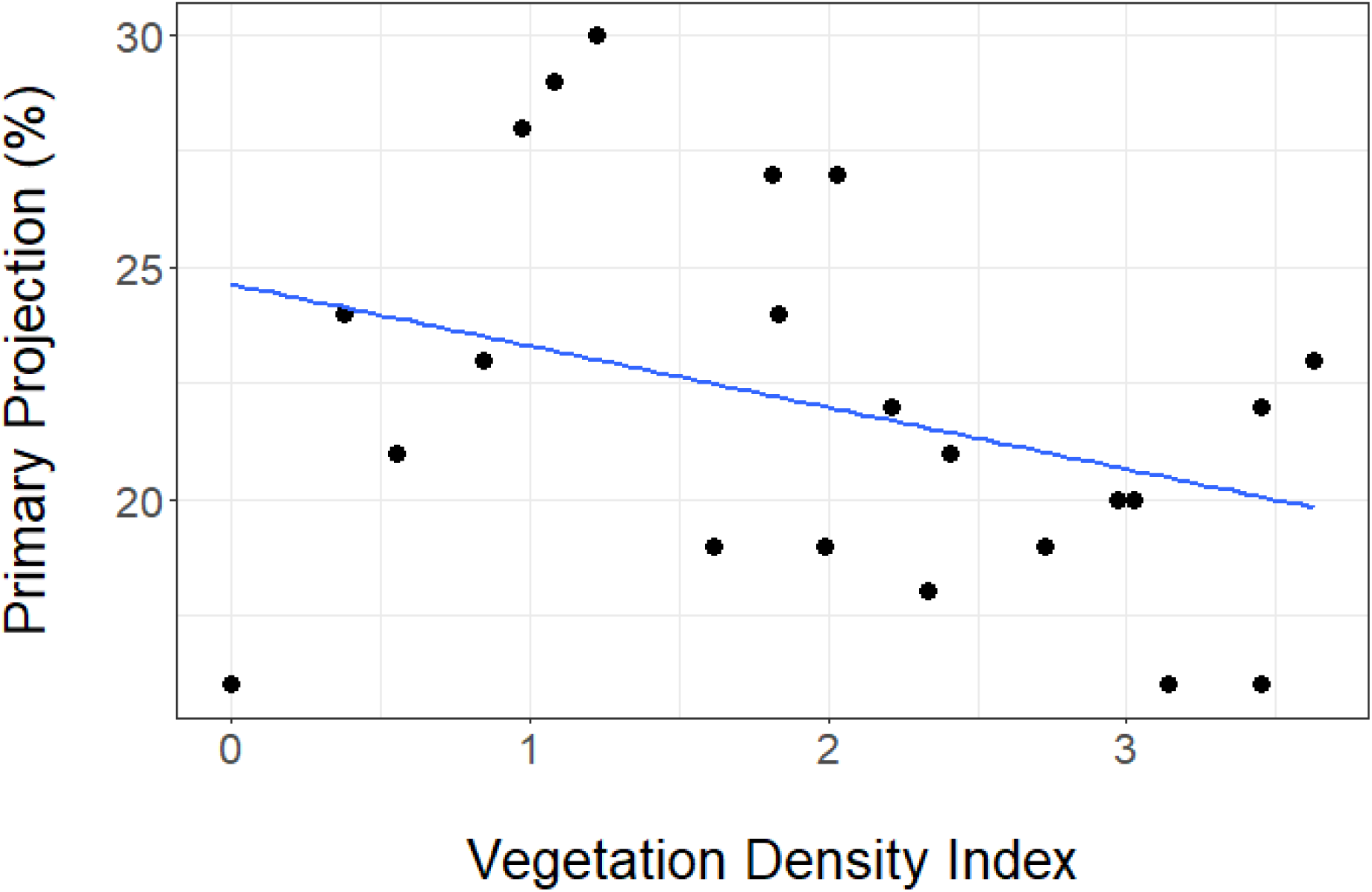
Wing pointedness in relation with vegetation density index. Each point denotes a species.

**Table 2.**
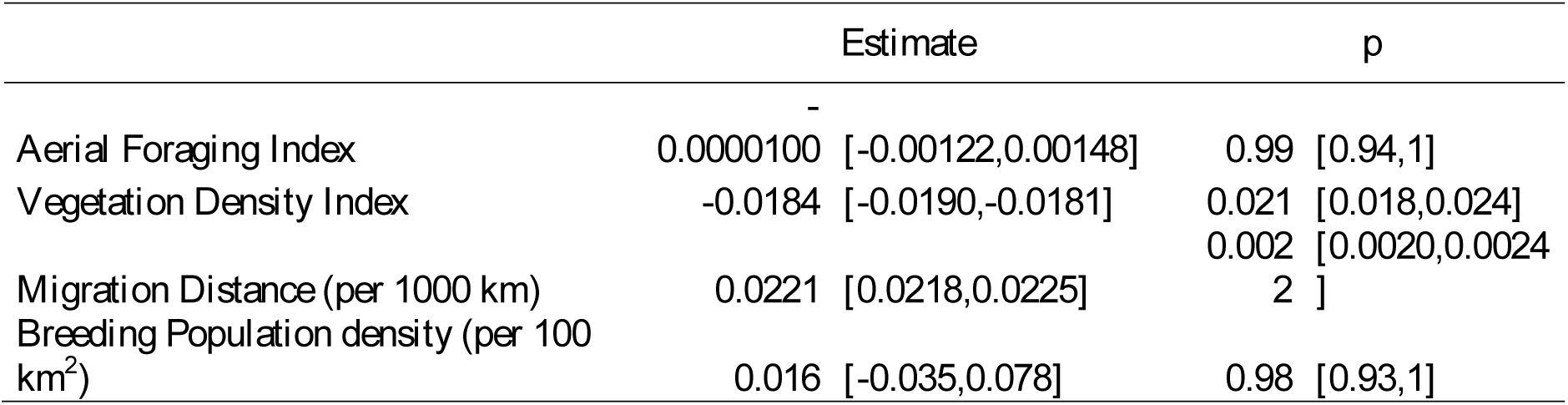
Means and ranges of GLS model estimates and p-values for 10,000 trees for relationship between wing pointedness, Aerial foraging Index, Vegetation Density Index, Migration Distance and Breeding Population Density (*N* = 22 species).

## Discussion

As we predicted, wing pointedness was greater in species that performed longer migration and smaller in species living in more densely vegetated environments. However, the amount of aerial foraging and population density had no explanatory power. The low variance of estimates between GLS models suggests that interspecific differences in wing shape and length (i.e. wing pointedness) were not greatly affected by uncertainty in phylogenetic relationships.

Since the early speculations by Palmer (1900) and Seebohm (1901) (cited by Mönkkönen (1995)), at least a dozen studies have provided evidence for more pointed or longer wings longer-distance migrants than conspecifics that migrate shorter distances (de La Hera et al. 2010; Rushing et al. 2014), or sedentary ones (Milá et al. 2008). Similar results were found in studies focusing on interspecific differences (Mönkkönen 1995; Minias et al. 2015). The present results, nearly half of which were obtained from north American warblers (Parulidae), are contrary to the earlier suggestions by Keast (1980) and Niemi (1985) that wing shape of those birds was unrelated to migration distance. Instead, this study adds to a growing body of evidence for a generally positive relationship between migratory behavior and wing attributes that facilitate sustained flight.

A notable exception to the wing shape – migration distance rule is the recent comparative analysis of eight swallow genera by Huber et al. (2016). They found a negative link between migration distance and wing pointedness, which they interpret as related to the swallows’ aerial foraging behavior which resembles their migratory flight, contrary to most other species whose wing pointedness vs. migration has been investigated so far. Earlier studies also pointed to the importance of foraging behavior as a determinant of wing pointedness, in bats (Baagøe 1987) and birds (Savile 1957; Marchetti et al. 1995; Vanhooydonck et al. 2009). Contrary to expectation, our results did not suggest a link between wing pointedness and aerial foraging *per se*, possibly because of insufficient variation in foraging methods among the 22 species, a lack of precision in our assessment of aerial foraging, or a combination of both. It would therefore be premature to rule out an ecomorphological effect of foraging behavior in itself on wing shape, at least in boreal songbirds.

Vegetation density has long been proposed as influencing wing shape, with longer-winged species found in more open habitats (Linsdale 1938). Rounded wings are often thought to allow better maneuverability, an asset in cluttered habitats (Norberg 1979). Furthermore, less pointed wings could facilitate predator escape on takeoff (Swaddle and Lockwood 2003; Fernández and Lank 2007), also advantageous mostly in dense cover. The present study is the first to provide evidence for a direct relationship between vegetation density and wing pointedness, but it would be premature to generalize to a broader set of species, because here we focused on a specific group of birds, i.e., small-sized forest birds predominantly foraging on standing vegetation or the ground.

Contrary to our expectation, there was no significant link between population density, which we assume reflects isolation (negatively), and wing pointedness. This could be due to insufficient precision in breeding population densities measures caused by the patchy records reported in BNA (Poole 2005) for most species. For example, those estimates do not distinguish between situations where a species would be found in dense clusters at the habitat patch scale, but scarce at the landscape scale, vs. uniformly scattered pairs throughout a landscape. Such differences could greatly affect the level of isolation experienced by individual members of the species. Furthermore, the population density of certain bird species may vary greatly from one region to another, as well as between wintering and nesting areas (Sherry and Holmes 1996). Unfortunately, there are few data on population densities in the wintering grounds. Regardless of data availability, it is perhaps unrealistic to expect that a species can be reduced to a single population density, because of the substantial geographic variation which is likely to occur.

Despite the latter caveats, the use of a comparative approach between different species of the same group still remains a useful approach to understand general patterns of habitat use in association to wing morphology. The accumulation of evidence on correlates of avian wing shape should not only lead to a better understanding of ecological differences among species, but opens the way for more incisive comparative analyses testing predictions about avian responses to ever-changing environments.

## Acknowledgements

Thanks to Vanessa Dufresne, Pierre-Alexandre Dumas, Jean-Michel Chabot and all volunteers, for their help in the field, and to the staff at Forêt Montmorency (Université Laval) for logistical support. This project was funded by an NSERC grant 170173 to AD, with authorization from Université Laval’s animal care committee (2013030-2).

